# Shaping information processing: the role of oscillatory dynamics in a working-memory task

**DOI:** 10.1101/2021.04.27.441620

**Authors:** Hesham A. ElShafei, Ying Joey Zhou, Saskia Haegens

## Abstract

Neural oscillations are thought to reflect low-level operations that can be employed for higher-level cognitive functions. Here, we investigated the role of brain rhythms in the 1–30 Hz range by recording MEG in human participants performing a visual delayed match-to-sample paradigm in which orientation or spatial frequency of sample and probe gratings had to be matched. A cue occurring before or after sample presentation indicated the to-be-matched feature. We demonstrate that alpha/beta power decrease tracks the presentation of the informative cue and indexes faster responses. Moreover, these faster responses coincided with an augmented phase alignment of slow oscillations, as well as phase-amplitude coupling between slow and fast oscillations. Importantly, stimulus decodability was boosted by both low alpha power and high beta power. In summary, we provide support for a comprehensive framework in which different rhythms play specific roles: slow rhythms control input sampling, while alpha (and beta) gates the information flow, beta recruits task-relevant circuits, and the timing of faster oscillations is controlled by slower ones.

**Significance statement:** Brain oscillations reflect low-level operations, building blocks, that control the flow of information through the brain. We propose and test a novel comprehensive framework in which slow oscillations control input sampling, alpha gates information flow, beta recruits task-relevant circuits, and the timing of faster oscillations is controlled by slower ones. We collected MEG data while participants performed a visual delayed match-to-sample task with pre- & retro-cues. Phase alignment of slow oscillations, governing input sampling, indexed faster responses. Alpha/beta power, gating information flow, boosted behavior & tracked informative cues. Low alpha (gating) & high beta (circuit-setup) power boosted signal information content. This is an essential step towards a more unified framework regarding the role of oscillatory dynamics in shaping information processing.

## 1 Introduction

Brain oscillations reflect rhythmic fluctuations of neuronal ensembles between states of low and high excitability (Bishop, 1932). Earlier research focused on investigating oscillations independently and linking them to high-level cognitive processes (e.g., theta to memory). However, this approach has been critiqued, as the exact role of each rhythm likely depends on an interplay between brain region, underlying neuronal substrate, and task context (Buzsaki, 2006; Wang, 2010). Our work is based on the hypothesis that oscillations reflect low-level mechanisms that can be flexibly employed to change the dynamics of neuronal populations, thereby providing the scaffolding for information processing. Thus, oscillations set the state of the system, either enhancing or attenuating activity (Klimesch et al., 2007), and inhibiting or facilitating network formation (Singer, 1999; Varela et al., 2001). Here, we focus on oscillations in the delta-to-theta, alpha, and beta bands in the context of working-memory.

Slow rhythms in the delta-to-theta bands have been proposed to govern the sampling of our surroundings by providing alternating phases of high and low neural excitability and thereby phases of high and low perceptual sensitivity (Fiebelkorn & Kastner, 2019; Helfrich et al., 2019; Herbst & Landau, 2016; VanRullen, 2016). Accordingly, sensory processing (and subsequent behavior) depends on the temporal coincidence of task-relevant targets with the oscillatory phase; i.e., when the phase with high excitability coincides with target occurrence, neural and behavioral responses are enhanced (Busch et al., 2009; Dugué et al., 2015; Fiebelkorn et al., 2013; Henry et al., 2016; Landau & Fries, 2012; VanRullen et al., 2011; Zion Golumbic et al., 2013). It has been shown that slow oscillations also control the timing of faster oscillations through phase-amplitude coupling (Canolty et al., 2010; Meij et al., 2012; Sauseng et al., 2019). However, it remains unclear how these cross-frequency interactions shape sensory processing and subsequent behavior. Here, we will investigate how the phase of slow rhythms, as well as coupling between slow and faster rhythms, correlates with behavior.

The alpha rhythm is thought to reflect functional inhibition (Klimesch et al., 2007), such that increased alpha power suppresses processing in task-irrelevant regions and networks (i.e., increased alpha reflects reduced neural excitability), while decreased alpha facilitates processing in task-relevant ones, effectively gating the information flow (Jensen & Mazaheri, 2010). Accordingly, it has been demonstrated that working-memory performance correlates positively with alpha activity in task-irrelevant (Haegens et al., 2010; Yu et al., 2017) and negatively in task-relevant regions (Jiang et al., 2015; van Ede et al., 2017), respectively. However, it remains unclear how alpha gating modulates internal sensory representations (as indexed by decoding performance), with studies reporting positive (i.e., decoding performance increases as alpha power increases; Kayser et al., 2016), negative (Barne et al., 2020), and null relationships (Griffiths et al., 2019). Here we will examine how alpha power correlates with working-memory performance as well as the decodability of task features.

While the beta rhythm was traditionally regarded as a somatomotor rhythm (Pfurtscheller & Lopes da Silva, 1999), recent evidence suggests that beta is involved in top-down processing (Buschman & Miller, 2007) and long-range communication (Engel & Fries, 2010). However, it is unclear onto which low-level mechanisms beta oscillations map. One proposal is that beta, like alpha, gates information processing through inhibition (Lundqvist et al., 2016; Miller et al., 2018). A more recent proposal is that beta facilitates the endogenous (re)activation of cortical content representations, e.g., during working memory (Spitzer & Haegens, 2017). In this framework, beta-band synchronization supports the endogenous transition from latent to active cortical representations during working memory, thereby providing a readout of the information held in memory. This proposal builds primarily upon evidence from non-human primate research (Bahmani et al., 2018; Haegens et al., 2017; Salazar et al., 2012) and is yet to be explicitly tested in the human brain (Herding et al., 2016). Here we will test the involvement of beta oscillations in circuit (re)activation, the possibility of the existence of multiple beta(s), and how these tentative different betas correlate with behavior.

Combined, we propose a framework of oscillations as building blocks, in which delta-to-theta rhythms sample task-relevant input, while the alpha rhythm suppresses irrelevant input. Crucially, the beta rhythm recruits task-relevant circuits that maintain information in working memory. Finally, the timing of faster oscillations is controlled by slower rhythms via phase-amplitude coupling. Critically, we propose that oscillations (separately and combined) orchestrate the functional architecture of information processing and thereby shape behavior. In order to elucidate unsettled oscillatory characteristics (including the relationship of alpha to sensory representations and phase-amplitude coupling to behavior), we recorded MEG in healthy participants preforming a visual delayed match-to-sample working-memory paradigm with pre- and retro-cueing.

## 2 Materials & Methods

### 2.1 Participants

Participants were 33 healthy right-handed adults (21 female, 12 male; mean age 24.6 years; range 20–33) without neurological or psychiatric disorders, who reported normal hearing and normal or corrected-to-normal vision. The study was approved by the local ethics committee (CMO 2014/288 “Imaging Human Cognition”) and in accordance with the Declaration of Helsinki. Participants gave written informed consent and were remunerated for their participation.

### 2.2 Visual working-memory task

We used a single-item delayed match-to-sample working-memory task where participants were instructed to compare sample and probe stimuli and indicate whether the cued feature was the same or different between them (Figure 1). Each trial contained four main events: a visual cue followed by a visual sample, a second visual cue, and a probe. All events were presented at a fixed stimulus onset asynchrony (SOA) of 1.5 s. The inter-trial interval (ITI) duration was jittered between 2 and 2.4 s.

**Figure 1.**
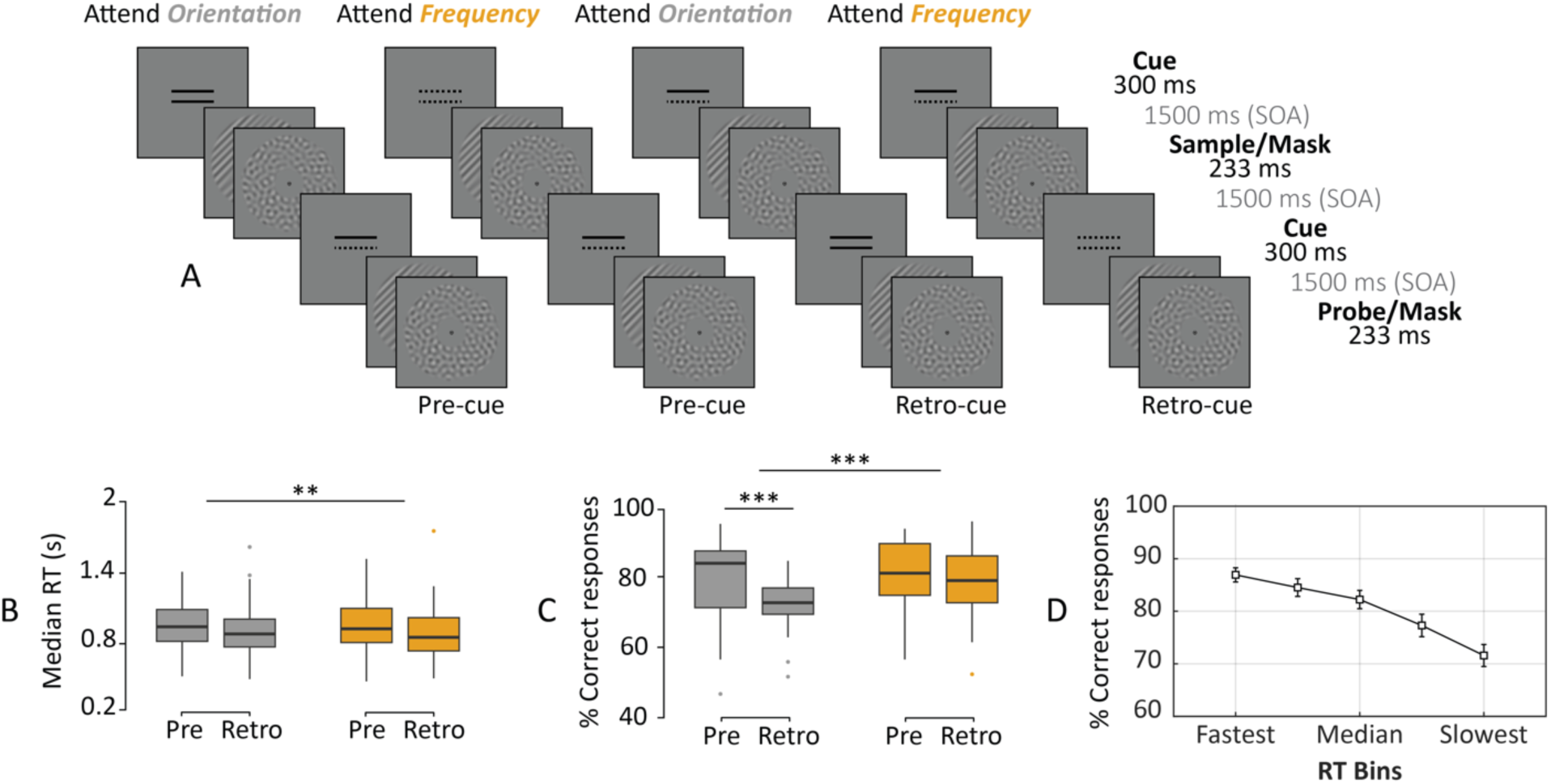
Paradigm and behavioral results. [A] Schematic of the four trial types: attend orientation (two solid lines) or attend frequency (two dotted lines) for pre- and retro-cue conditions (first vs. second cue is informative; uninformative cues consist of a solid and dotted line). [B] Participants were slower and [C] more accurate matching the spatial frequency (gray bars) of the visual gratings compared to orientation (orange). This effect was mainly driven by the pre-cue condition. Within each boxplot, the horizontal line represents the median, the box delineates the area between the first and third quartiles (interquartile range). [D] Average accuracy across RT bins. Error bars represent standard error of the mean. ** *P* < 0.01, *** *P* < 0.001.

We used a two (relevant feature: orientation vs. frequency) by two (informative cue: pre- vs. retro-cue) factorial design. Participants were cued to compare sample and probe orientation in half of the trials, and spatial frequency in the other half. They were informed about the relevant feature by an informative cue, which occurred either before (pre; 50%) or after (retro; 50%) the sample presentation. These four trial types were randomly interleaved such that within each block participants performed all task conditions. Participants gave their response (“same” or “different”) by pressing a button with their right index or middle finger. The response-button mapping changed on a block-by-block basis in order to avoid motor preparation confounds in our decision-related analyses.

Participants first performed a few short practice sequences of the task (20 trials each) in which feedback was provided on a trial-by-trial basis, with green and red fixation dots for correct and incorrect responses, respectively. During the main experiment, we recorded MEG while participants performed 8 blocks of 64 trials each, in which feedback was provided on a block-by-block basis. The full recording session lasted around 90 minutes.

### 2.3 Stimuli

A bull’s eye (outer black ring = 0.5° × 0.5° degree of visual angle (dva), innermost black dot = 0.25° × 0.25° dva) was presented at the center of the screen as the fixation point. Participants were instructed to always maintain fixation, and not to blink during the presentation of the stimuli. Each trial started with a cue presented for 300 ms. Next, a sample consisting of an oriented grating (Michelson contrast: 40%, spatial frequency: 1 cycle per °, orientation: 45° clockwise or counterclockwise relative to vertical, randomized spatial phase) and a backward mask (a bandpass-filtered noise patch with matched contrast and spatial frequency as the grating) were presented sequentially for 230 ms. Both the grating and the mask were shown in an annulus (inner radius = 1.5°, outer radius = 7.5°, contrast of the stimuli decreased linearly to 0 over the outer and inner 0.5° radius of the annulus) around the central fixation (Figure 1A). After an SOA of 1.5 s, a second cue was presented for 300 ms. The trial presentation finished with the probe presentation, which, similar to the sample, consisted of an oriented grating and a backward mask. The probe’s cued feature matched with that of the sample grating in 50% of the trials. The experiment was programmed with PsychtoolBox (Brainard, 1997) in Matlab (Mathworks, Inc.).

### 2.4 Data acquisition

Stimuli were displayed on a semitranslucent screen (1920 × 1080-pixel resolution, 120-Hz refresh rate) back-projected by a PROpixx projector (VPixx Technologies) during MEG recordings. Whole-head MEG data were acquired at a sampling frequency of 1200 Hz with a 275-channel MEG system with axial gradiometers (CTF MEG Systems, VSM MedTech Ltd.) in a magnetically shielded room. Six permanently faulty channels were disabled during the recordings, leaving 269 recorded MEG channels. Three fiducial coils were placed at the participant’s nasion and both ear canals, to provide online monitoring of participant’s head position (Stolk et al., 2013) and offline anatomical landmarks for co-registration. Eye position was recorded using an eye tracker (EyeLink, SR Research Ltd.). Upon completion of the MEG session, participant’s head shape and the location of the three fiducial coils were digitized using a Polhemus 3D tracking device (Polhemus, Colchester, Vermont, United States). Anatomical T1-weighted MRIs were obtained during a separate session. To improve co-registration of the MRIs and MEG data, earplugs with a drop of Vitamin E were placed at participant’s ear canals during MRI acquisition. These anatomical scans were used for source reconstruction of the MEG signals.

### 2.5 Behavioral analysis

The influence of cue condition (two levels: pre- and retro-cue) and feature (two levels: orientation and spatial frequency) on median reaction time (RT; including correct responses only) and accuracy (percentage of correct responses) was tested using a linear mixed-effects model using the lme4 package (Bates et al., 2015) for R (Team, 2014). For post-hoc analysis we used the Lsmean package (Searle et al., 1980) where p-values were considered as significant at *p*<0.05 and adjusted for the number of comparisons performed (Tukey method).

### 2.6 MEG pre-processing

MEG data were preprocessed offline and analyzed using the FieldTrip toolbox (Oostenveld et al., 2011) and custom-built MATLAB scripts. The MEG signal was epoched based on the onset of the first cue (t= -1 to 7s). The data were downsampled to a sampling frequency of 300 Hz, after applying a notch filter to remove line noise and harmonics (50, 100, and 150 Hz). Bad channels and trials were rejected via visual inspection before independent component analysis (Jung et al., 2001) was applied. Subsequently, components representing eye-related and heart-related artefacts were projected out of the data (on average, four components were removed per participant). Finally, outlier trials of extreme variance were removed. This resulted in an average of 393 (± 9 SEM) trials and 268 channels per participant for the reported analyses.

### 2.7 Event-related fields

Single-trial data were baseline corrected (t = −0.1 to 0 s) before calculation of first-cue and grating locked event-related fields (ERFs). In addition, a planar representation of the MEG field distribution was calculated from the averaged data (ERFs) using the nearest-neighbor method. This transformation makes interpretation of the sensor level data easier as the signal amplitude is typically maximal above a source.

### 2.8 Spectral analysis

Our analysis aimed to highlight the (1) differential impacts of pre- and retro-cueing on the oscillatory dynamics of interest, and (2) the behavioral relevance of these dynamics. For both these contrasts, our spectral measures included delta-to-theta inter-trial phase coherence, alpha power, and beta power. For the cueing contrast, spectral measures were compared between pre- and retro-cue trials (including correct response trials only). For the behavioral contrast, trials from all conditions (including both correct and incorrect response trials) were pooled together and divided into five quantile bins (approx. 80 trials per bin) according to RT. Spectral measures were contrasted between the slowest and fastest RT bins, to maximize statistical power.

For the 20 MEG channels displaying the maximal post-grating ERFs (channels selected individually per participant), we extracted 1-s pre-stimulus windows (t = −1 to 0 s), multiplied these with a Hanning taper, and computed power spectra (1–30 Hz; 1-Hz resolution) using a fast Fourier transform (FFT) approach. In order to determine the individual alpha peak frequency, we detected the highest local maximum within the 7–14 Hz band. In order to detect the beta peak frequency, linear regression (least-squares fit) was used to fit a linear model to the log-transformed spectrum in the beta range in order to compensate for the 1/f effect. The fitted linear trend was then subtracted from the spectrum, allowing for a more reliable beta peak frequency estimate within the 15–30 Hz band (Haegens et al., 2014).

We identified distinct alpha peak frequencies for all participants (mean = 9.9 Hz ± 0.21 SEM; Figure 2), while for beta we identified peaks for 18 participants (mean = 20.8 Hz ± 0.37 SEM) and assigned the mean beta peak for the remainder of the participants (N=15). It is important to note that our peak estimations did not significantly vary when removing the 1/f background noise (p = 0.1 for alpha, p = 0.3 for beta). In order to create peak-centered signals, we computed TFRs of the power spectra for the full trials per experimental condition. To this end, we used an adaptive sliding time window of five cycles length per frequency (Δt = 5/f; 20-ms step size), and estimated power after applying a Hanning taper. We created alpha-peak (individualized) and beta-peak (individualized) centered time-resolved signals with spectral bandwidths of 2 Hz and 4 Hz, respectively. Power normalization was performed relative to a baseline (t = -0.4 to -0.2 s) for the cueing contrast and relative to the average of all RT bins for the RT contrast.

**Figure 2.**
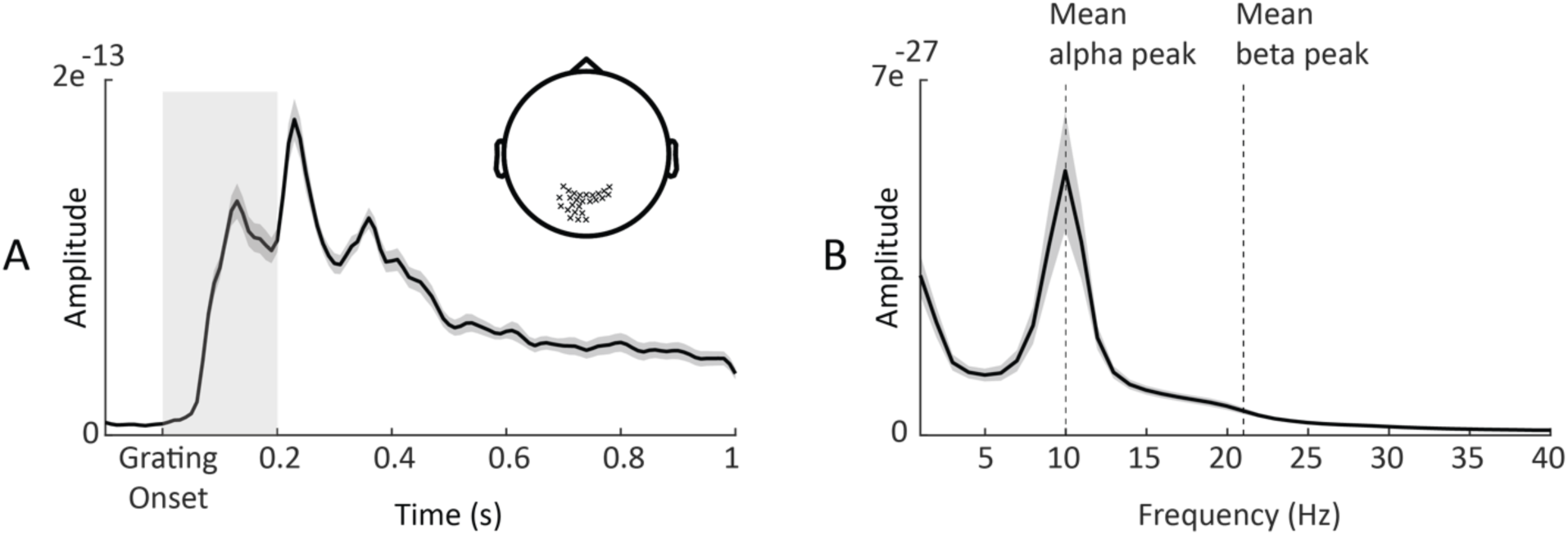
Alpha and beta peak detection. [A] Grand-averaged ERF of sensors (highlighted in sensor space) displaying maximal post-grating ERFs within 200 ms following the grating onset. Gray shaded area around the curve denotes between-participants standard error. Light gray shaded box highlights the time period used to select channels with maximal post-grating ERFs. [B] Power spectra (averaged over same sensors marked in A; t = -1 to 0 s relative to first cue onset) showing alpha and beta peaks.

We estimated inter-trial phase coherence (Tallon-Baudry et al., 1996) across trials, for each time point, frequency and sensor, for each RT bin separately. This measure reflects the degree to which the phase of each frequency is aligned across trials. We averaged this measure between 1 and 6 Hz in order to reduce data dimensionality. In addition, we used the Tensorpac toolbox (Combrisson et al., 2020) to compute a time-resolved measure of phase-amplitude coupling (PAC) on source level. This algorithm measures PAC across trials (rather than across time).

### 2.9 Statistical analysis

In order to investigate whether differences between conditions of interest (cueing contrast: pre- vs. retro-cue; RT contrast: slowest vs. fastest RT bin) were statistically significant, we used nonparametric cluster-based permutation analysis (Maris & Oostenveld, 2007). In brief, this test first calculates paired t-tests for each sensor at each time and/or frequency point, which are then thresholded at *p* < 0.05 and clustered on the basis of temporal, spatial and/or spectral adjacency. The sum of t-values within each cluster is retained, and the procedure is repeated 1000 times on permuted data in which the condition assignment within each individual is randomized. On each permutation, the maximum sum is retained. Across all permutations, this yields a distribution of 1000 maximum cluster values. From this distribution, the probability of each empirically observed cluster statistic can be derived (evaluated at alpha = 0.05).

### 2.10 Source reconstruction

For each participant, an anatomically realistic single-shell headmodel based on individual T1- weighted anatomical images was generated (Nolte, 2003). A grid with 0.5-cm resolution was created using an MNI template onto which the brain volume of each participant was morphed using non-linear transformation. For each grid point, leadfields were computed with a reduced rank, which removes the sensitivity to the direction perpendicular to the surface of the volume conduction model. This procedure ensures that each grid point represents the same anatomical location across all participants.

In order to visualize the sources of our sensor-level spectrotemporal (power and ITC) and temporal effects (ERFs), we utilized the frequency-domain adaptive spatial technique of dynamical imaging of coherent sources (DICS; Gross et al., 2001) and the linearly constrained minimum variance beamformer (LCMV; Van Veen et al., 1997), respectively. Data from all conditions of interest were concatenated in order to compute the cross-spectral density (CSD) matrices (t= -1 to 2 s; multitaper method (Mitra & Pesaran, 1999)) and the covariance matrices of the averaged single trials (t= -0.6 to 5.5 s; lambda 5%) for the DICS and LCMV beamformer, respectively. Leadfields for all grid points along with the CSD/covariance matrices were used to compute a common spatial filter (i.e., common for all trials and conditions) that was used to estimate the spatial distribution of power/amplitude for our windows of interest. The source orientation was fixed to the dipole orientation with the highest strength.

Finally, the PAC analysis was performed on source level (i.e., virtual channels), for which the source space was subdivided into 22 anatomically defined brain parcels, including the intraparietal sulci, superior parietal lobes, frontal eye fields, middle frontal gyri and the temporoparietal junction (Wallis et al., 2015). Sensor-level time-series data were multiplied by spatial filters (constructed using LCMV beamformer) in order to obtain the time-resolved activity in each virtual channel.

### 2.11 Multivariate pattern analysis

To investigate how oscillatory dynamics influence the information content of neural activity, we were interested in decoding two task features: (1) informative cue instruction (attend orientation or spatial frequency), and (2) sample (i.e., the first grating) properties (orientation: clockwise or counterclockwise; spatial frequency: low or high). For all multivariate pattern analysis (MVPA), we used MNE Python (Gramfort et al., 2013) and Scikit-learn (Pedregosa et al., 2011). We implemented a 4-fold cross-validation procedure within each participant. For each timepoint, we trained a logistic regression classifier (L2-regularized) on three folds and tested on the left-out fold. The analysis was shifted over time on a sample-by-sample basis. The input signal was the broadband signal lowpass filtered at 35 Hz (Gwilliams & King, 2020; King et al., 2016). We operationalized the decoding accuracy as area under the curve (AUC) by evaluating the similarity between the true label categories of the test set and the probabilistic class labels (normalized distance from the fit hyperplane: “predict_proba” in scikit-learn) of the same trials. We computed the AUC under the null hypothesis by randomly shuffling the label categories and obtained a p-value as the proportion of the null AUC estimates that exceeded the true AUC (evaluated at alpha = 0.05).

We aimed to first demonstrate the decodability of the aforementioned features from the broadband time-domain signal. We used trials from all conditions and tested decoding accuracy against chance level (i.e., AUC = 0.5 for binary classification) using nonparametric cluster-based permutation analysis (as described above). All task features could be decoded from the MEG signal: informative cue instruction (attend to frequency or orientation), grating orientation (clockwise or counterclockwise) and spatial frequency (low or high). For cue identity, group-level decoding performance peaked around 150 ms after the onset of the first/second cue for the pre-/retro-cue conditions, respectively (Figure 3AB), while for grating properties, group-level decoding performance (for orientation and spatial frequency) peaked around 100 ms (Figure 3CD). It is important to note that the decodability of the grating features correlated negatively with reaction times (r = -0.5, p < 0.006) and positively with accuracy (r = 0.3, p = 0.042). This provides evidence that the decoded information reflects internal representation that are behaviorally relevant (Ritchie et al., 2019).

Next, we asked whether oscillatory state shapes neural stimulus representations as reflected by decoding performance. To test this, we evaluated how decoder accuracy was related to trial-by-trial fluctuations of prestimulus power. We used the classification procedure explained above, replacing the k-fold cross validation with a leave-one-out cross-validation (LOOCV) procedure. In LOOCV, the classifier is fit to all trials but one, evaluating model performance on the remaining “left-out” trial as a single-item test set. This is advantageous because: (1) it allows a maximal amount of data to be used for training, thus reducing noise in the model fit; (2) it provides an unbiased single-trial decoding estimate, which can be analyzed by a binning approach. For each timepoint and test trial, we computed the probabilistic estimates of the logistic regression for our feature of interest (cue information) and then grouped these results into five bins relative to occipital sensor theta, alpha and beta power. Afterwards, for each frequency band separately, the time-resolved AUC signal was contrasted between the bins of lowest and highest power using nonparametric cluster-based permutation analysis.

Finally, to investigate the evolution of sample representations, we utilized the temporal generalization method (King & Dehaene, 2014). For this analysis, each classifier was trained on time *T* and tested on its ability to predict a given trial at time *T’*. This method estimates the similarity of the coding pattern at T and *T’,* and thus the stability of the neural representation. In order to facilitate the interpretation of our results, we averaged the AUC for training times within the first 0.3 s following the sample onset. This was done separately for each bin and each band, followed by contrasting the averaged temporal generalization matrices between the bins of lowest and highest power using nonparametric cluster-based permutation analysis.

### 2.12 Data and code availability

All data and code used for stimulus presentation and analysis are available online at the Donders Repository at https://doi.org/10.34973/tqy5-mh37.

## 3 Results

### 3.1 Behavioral results

In order to test our framework, we recorded MEG data from 33 healthy participants who performed a delayed match-to-sample working-memory paradigm where they evaluated orientation or spatial frequency of two visual gratings. The to-be-matched feature was indicated by a visual cue occurring either before (pre-cue) or after (retro-cue) sample presentation.

Overall, participants were faster matching the orientation than the spatial frequency of the gratings (F(1,32) = 11.2, p = 0.001; Figure 1B). There was no significant difference in RT between cue conditions (F(1,32) = 2.7, p = 0.07), nor a significant interaction between feature and cue conditions (F(1,128) = 12.7, p = 0.66). In terms of accuracy, participants were better at matching spatial frequency than orientation (F(1,128) = 13.4, p < 0.001; Figure 1C). There was no significant difference in accuracy between cue conditions (F(1,128) = 2.7, p = 0.10), though there was a significant interaction between the cue and feature conditions (F(1,128) = 12.7, p < 0.001): when matching the orientation of gratings, participants were better in the pre-cue than the retro-cue condition (t(32) = -3.7, p < 0.001), whereas when matching the spatial frequency of gratings, participants performed similarly in both cue conditions (t(32) = 1.35, p = 0.17).

Note that the absence of main cue effects demonstrates that task difficulty (performance) was comparable across our conditions of interest (i.e., pre- and retro-cue) and therefore does not constitute a confound in our subsequent analyses. Furthermore, accuracy linearly decreased with increasing RT (R^2^ = 0.21, p < 0.001; Figure 1D), suggesting no speed-accuracy tradeoff.

### 3.2 Impact of cueing on oscillatory dynamics

In order to establish the proposed low-level mechanisms, we first investigated the differential impacts of pre- and retro-cueing on oscillatory delta-to-theta phase, alpha power, and beta power. The critical difference between the pre- and retro-cue conditions is the time when participants are informed about the task-(ir)relevant feature. If alpha/beta decrease is critical for the inhibition of task-irrelevant features, we expect alpha/beta power decrease to be contingent upon the informative cues, after which selection (i.e., inhibition of the task-irrelevant feature and/or enhancement of the relevant one) is possible. Similarly, if delta-to-theta phase plays a role in input sampling, we would expect an increase in phase alignment following the informative cues.

Indeed, we found significant differences between pre- and retro-cue conditions for all our frequency bands of interest (Figure 4). Following the first cue, alpha and beta power decrease was stronger for the pre-cue condition (in which the first cue was informative) in comparison to the retro-cue condition (Figure 4CD; cluster-based permutation test, p = 0.002 and 0.003; t = 0.35 to 1 s and 0.34 to 0.77 s; centered on occipitoparietal and temporal sensors). Differences between pre- and retro-cue conditions were mainly localized to left and right occipital cortices, and right middle and superior temporal gyri.

**Figure 3.**
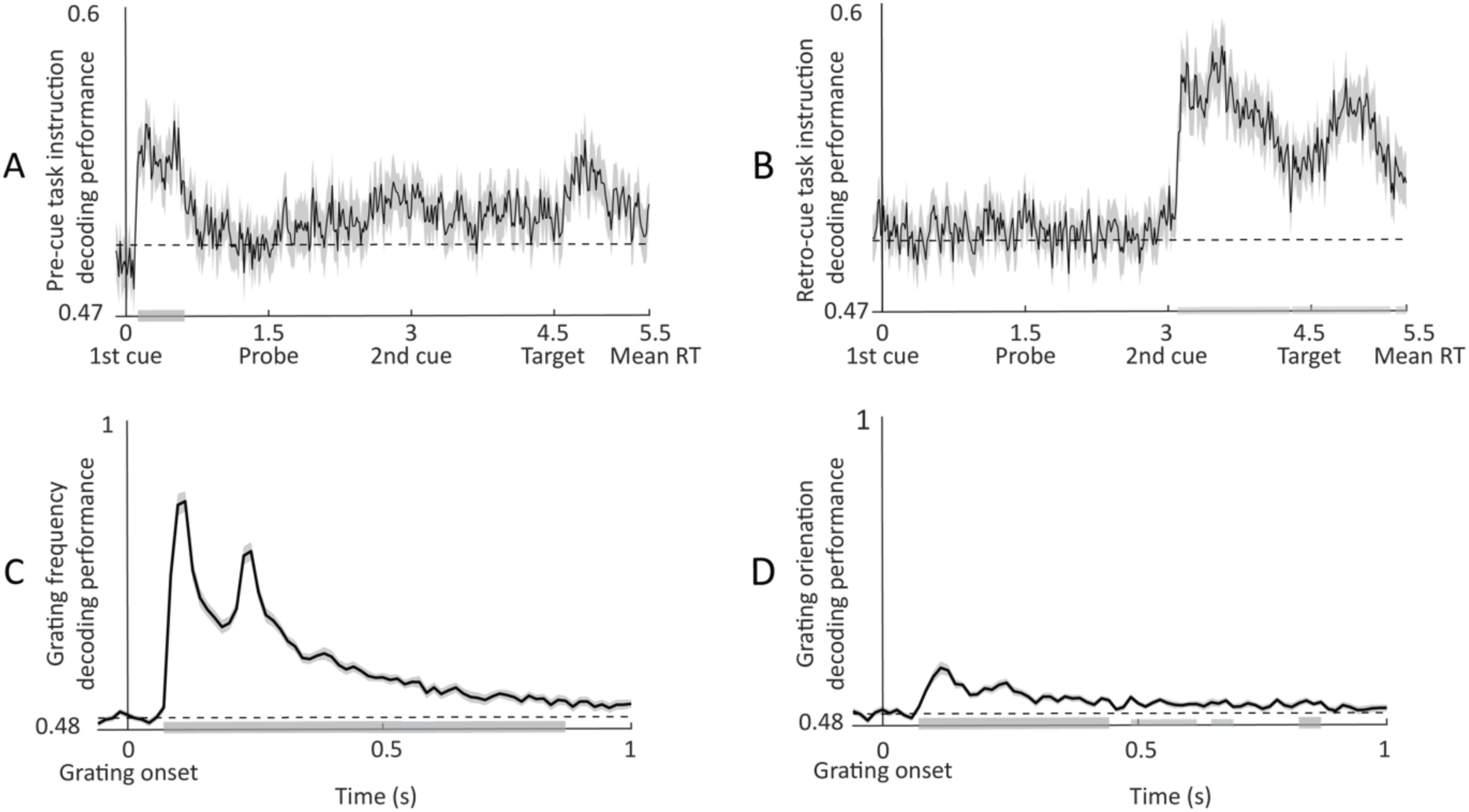
Decodability of task features. [A] Temporal sample-by-sample decoding of cue features (attend frequency vs. attend orientation) for pre- and [B] retro-cues. [C] Decoding performance for grating spatial frequency (low vs. high) and [D] grating orientation (clockwise vs. counterclockwise). Gray bars indicate significance of decoding accuracy (t-test vs. chance).

**Figure 4.**
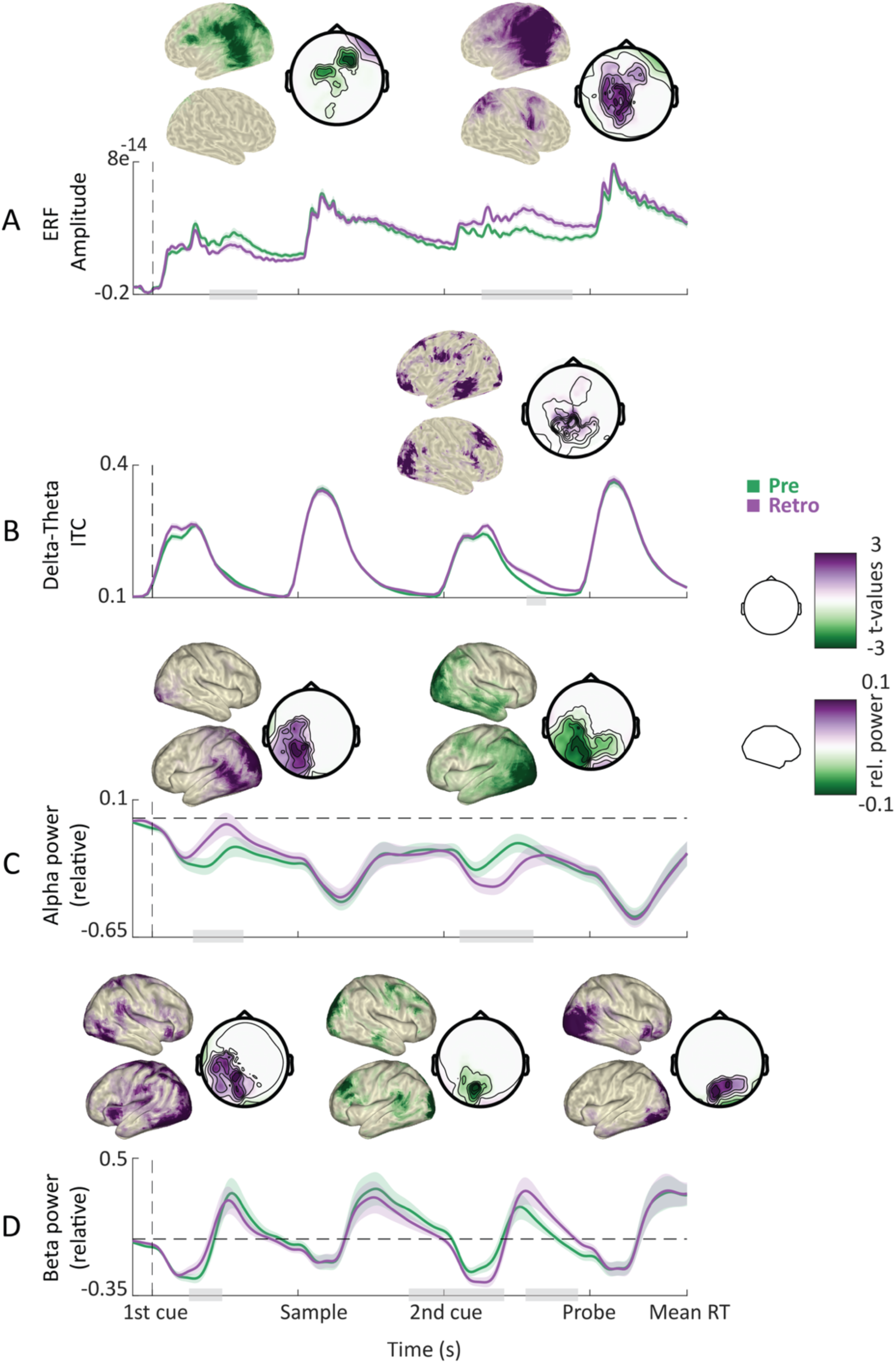
Cue-related power modulations. [A] Grand average ERFs for the pre- (green) and retro-cue conditions (purple). Shaded areas denote between-participants standard error. Gray bars indicate significant differences between conditions. Topographies show statistical and power distributions of these significant differences in sensor and source space, respectively. [B] Inter-trial phase coherence (averaged between 1 and 6 Hz) for the pre- and retro-cue conditions. [C] Time course of oscillatory power averaged within the alpha band. [D] Same as C but for the beta band. For all time courses, all sensors displaying significant differences as highlighted by the topographies were included in the plot.

This pattern was reversed following the second cue, with alpha and beta power decrease being stronger for the retro-cue condition (in which the second cue was informative) in comparison to the pre-cue condition (Figure 4CD; p = 0.001; t = 3 to 4 s and 2.6 to 3.6 s; centered on occipitoparietal sensors). In addition, beta power decrease was stronger for the pre-cue vs. retro-cue condition just prior to the probe presentation (Figure 4D; p = 0.004; t = 3.8 to 4.4 s; centered on occipital sensors). Differences between pre- and retro-cue conditions were mainly localized to left and right occipital cortices and the precentral gyrus, with beta effects extending more towards the dorsolateral prefrontal cortex.

Furthermore, following the second cue, delta-to-theta inter-trial phase coherence was stronger for the retro-cue condition in comparison to the pre-cue condition (Figure 4B; p = 0.001; t = 3.6 to 4.1 s; centered on occipital, parietal and temporal sensors). Differences between pre- and retro-cue conditions were mainly localized to left and right occipital cortices, left inferior and middle temporal gyrus, and left and right prefrontal cortices.

To summarize, the informative cue (i.e., the first cue on pre-cue trials and the second cue on retro-cue trials) was followed by a stronger decrease in alpha and beta power, consistent with the functional inhibition account. In addition, the informative cue on retro-cue trials was followed by an increased phase alignment in the delta-to-theta band, consistent with a role in regulating input sampling.

### 3.3 Impact of oscillatory dynamics on behavioral performance

Next, we asked whether any of the observed oscillatory dynamics impacted behavioral performance. We expected that faster responses would coincide with (1) an increase in the phase alignment of slow rhythms, reflecting their role in input sampling and temporal prediction, (2) a decrease in occipital alpha/beta power, reflecting their gating role, and (3) increased phase-amplitude coupling between slow and fast rhythms, reflecting the temporal coordination of fast rhythms by slower ones. We binned trials into five bins based on RT and contrasted oscillatory dynamics between bins of the slowest and fastest RT to test these predictions.

In line with our predictions, faster responses were associated with weaker alpha and beta power before and during probe presentation (Figure 5CD; p = 0.002 and 0.001; t = 4.1 to 5.1 s; centered on occipitoparietal sensors). These effects were localized to occipital cortices and the left inferior and parietal cortices. Furthermore, faster responses were accompanied by an increase in ITC in delta-to-theta frequencies (Figure 5B; p < 0.001; t = 4.3 to 5.5 s; across widespread sensors). Differences were maximal in left and right post- and precentral gyri and lateral prefrontal cortices.

**Figure 5.**
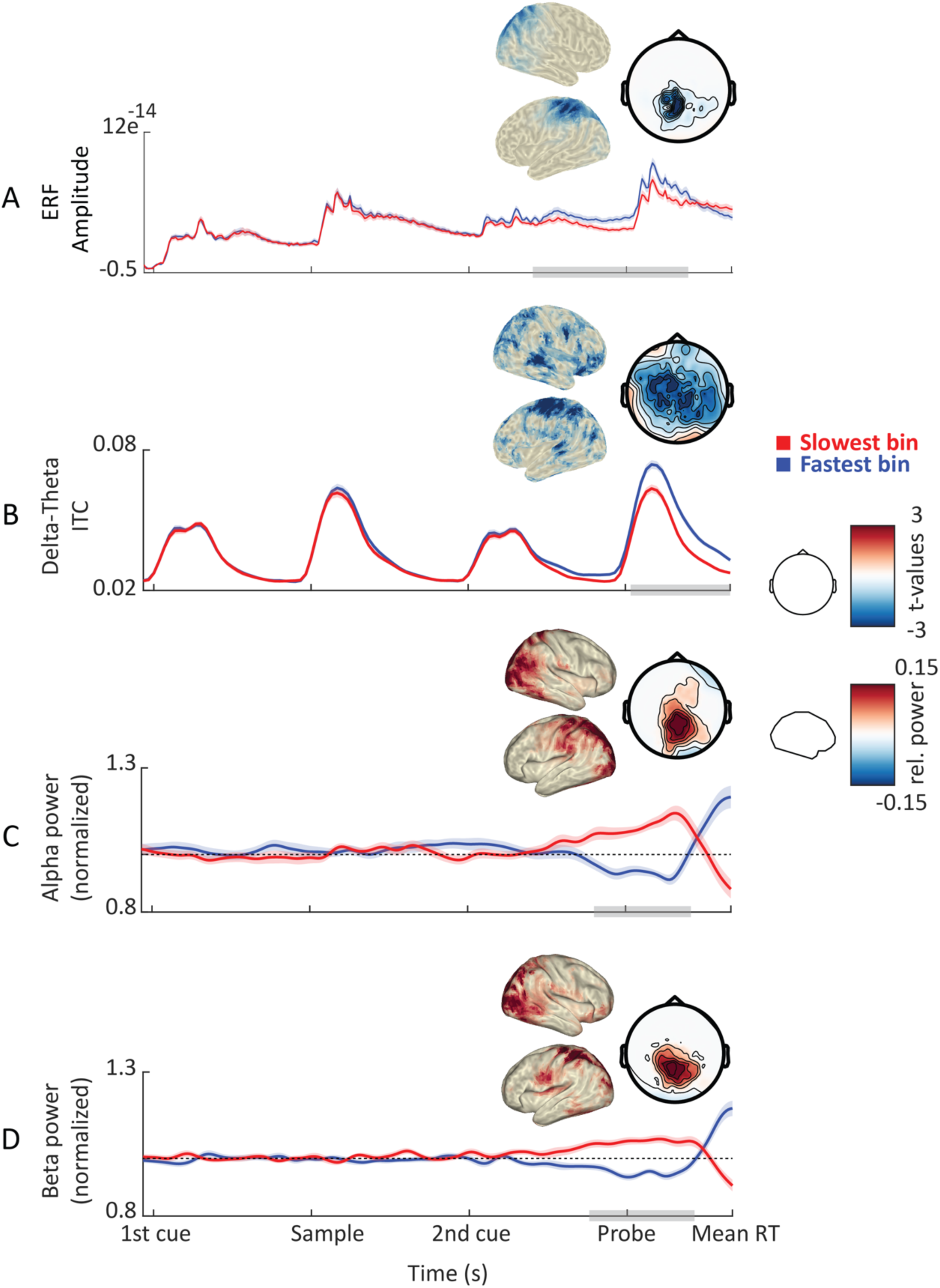
Correlates of behavioral performance: power and phase. [A] Grand average ERFs (occipital sensors) for the slowest (red) and fastest RT bins (blue). Shaded areas denote between-participants standard error. Gray bars indicate significant differences between conditions. Topographies show statistical and power distributions of these significant differences in sensor and source space, respectively. [B] Time course of inter-trial phase coherence (averaged between 1 and 6Hz) for the slowest and fastest RT bins. [C] Time course of oscillatory power averaged within the alpha band. Dashed lines represent mean power (i.e., normalized power = 1). [D] Same as C but for the beta band.

We found that the coupling between the phase of 1–3 Hz oscillations and the amplitude of 30– 35 Hz oscillations was stronger for fast vs. slow RT in the intraparietal sulcus (Figure 6C; p = 0.02; t = 0 to 5.5 s) and the superior parietal lobe (Figure 6D; p = 0.07; t = 0 to 3 s). Combined, these results indicate that response speed was influenced by the phase alignment of delta-to-theta oscillations as well as phase-amplitude coupling between slow and fast rhythms.

**Figure 6.**
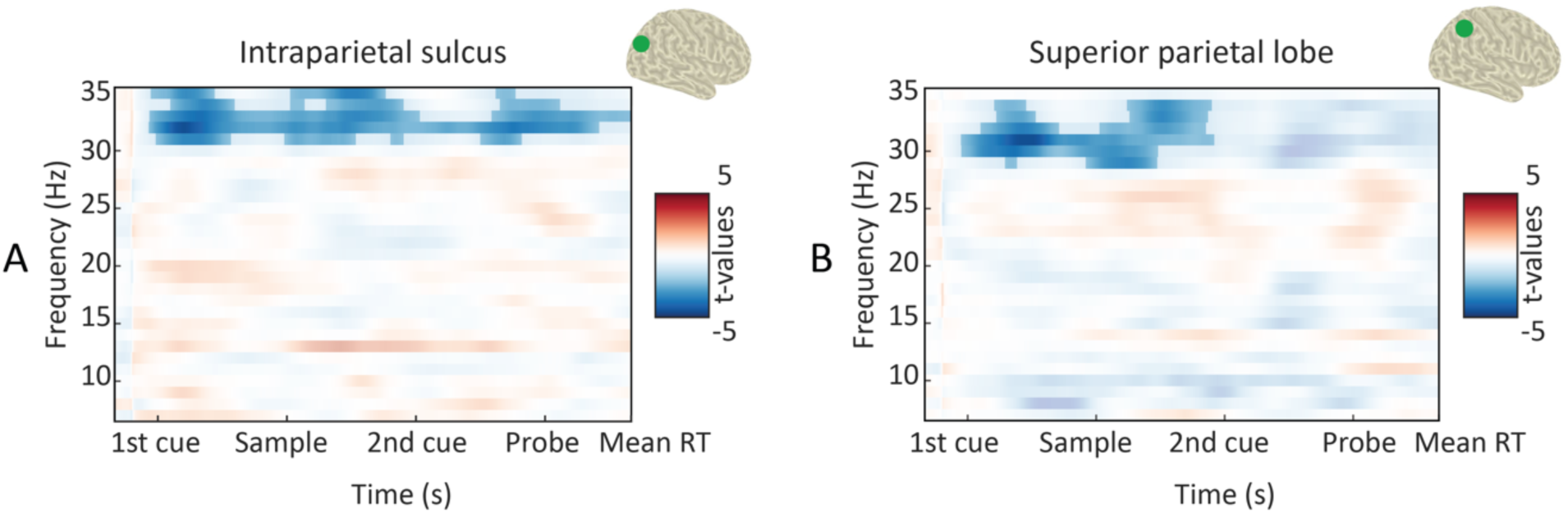
Correlates of behavioral performance: ITC and PAC. [A] TFR of statistical differences in PAC between the slowest and fastest RT bins (masked at p < 0.05) in the intraparietal sulcus (as highlighted in green on brain surface). Negative values (blue) indicate higher PAC for the fastest RT bin. [B] Same as A for the superior parietal lobe.

In sum, faster RT was preceded by (1) an increase in phase alignment and power of slow oscillations, consistent with a role in regulating input sampling, (2) a prominent occipitoparietal alpha/beta decrease, consistent with a gating-through-inhibition mechanism, and (3) an increased phase-amplitude coupling between slow and fast oscillations, consistent with a role of slow oscillations in temporal control of faster oscillations.

### 3.4 Disentangling oscillatory dynamics from evoked responses

To ensure that the ITC results (for the cueing and RT contrasts) did not stem from low-frequency evoked (phase-locked) responses, we ran additional (control) analyses. For the cueing contrasts, we compared ERFs between cue conditions and found that following the first cue, ERF amplitude was higher for the pre-cue compared to the retro-cue (Figure 4A; p < 0.001; t = 0.6 to 1.1 s; centered on centroparietal sensors). In addition, following the second cue, ERF amplitude was higher for the retro-cue (Figure 4A; p < 0.001; t = 3.4 to 4.5 s; widespread). Further, we compared the ERF and ITC differences between conditions at source level and found that ERF differences were more prominent in right pre- and post-central gyri and prefrontal cortex (Figure 7A).

**Figure 7.**
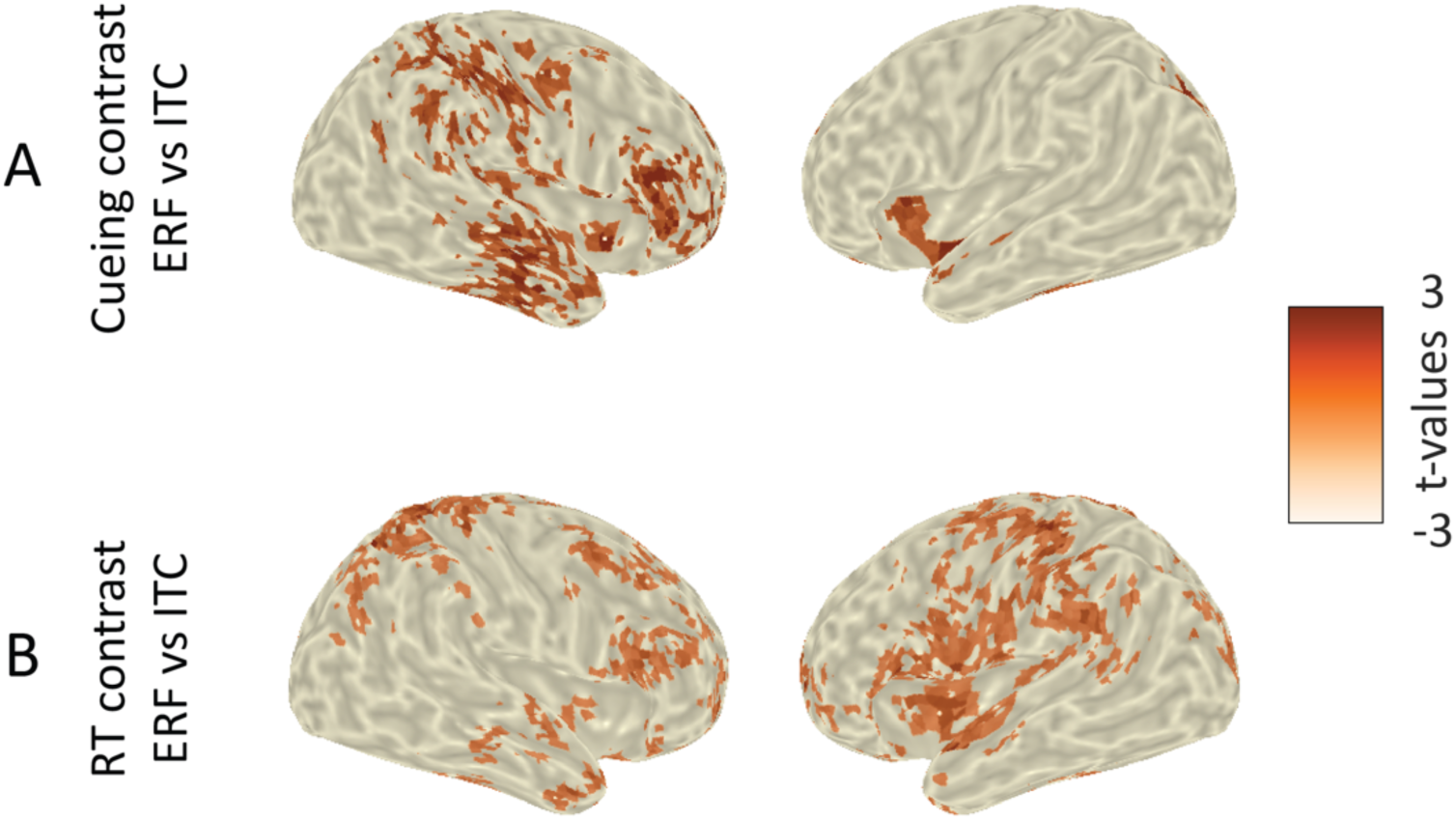
Correlates of behavioral performance: ERFs. [A] Source level topographies of the statistical differences between ERF and ITC for the cueing contrast (pre- versus retro-cue; t=3.6 to 4.1 s; masked at p < 0.05). [B] Same as A for the RT contrast (slow versus fast RT bin; t= 4.4 to 5.1 s).

For the RT contrast, we compared ERFs between RT bins, and found that ERF amplitude was higher for the fastest compared to the slowest RT bin (Figure 5A; p < 0.001; p < 0.05; t = 3.7 to 5.5 s; centered on occipitoparietal sensors). Further, we compared the ERF and ITC differences between RT bins at source level and found that ERF differences were more prominent in left and right pre- and post-central gyri and prefrontal cortex (Figure 7B). The significant differences between the localization of the ERF and ITC effects suggest that these effects (ERF and ITC) might reflect two separate phenomena.

Finally, we repeated the ITC analyses after subtraction of the evoked response from single-trial data. The results were similar to those of our initial analysis: for the cueing contrast, in the retro-cue condition, the presentation of the second cue was accompanied by an increase in ITC (p < 0.001; t = 3.5 to 4.3 s; widespread). For the RT contrast, faster responses were accompanied by an increase in ITC (p < 0.001; t = 4.7 to 5.4 s; widespread). The persistence of effects after the subtraction of evoked responses indicates that the low-frequency modulations cannot be fully explained by evoked activity.

### 3.5 Impact of oscillatory dynamics on decoding performance

Finally, we investigated how oscillatory dynamics influence the information content of neural activity patterns. We expected this content, indexed by classification accuracy, to be boosted by a decrease in alpha power (i.e., release of inhibition) and an increase in beta power (reflecting circuit recruitment). We trained neural classifiers to decode two task features: (1) informative cue instruction (attend orientation or spatial frequency), and (2) grating properties (orientation: clockwise or counterclockwise; spatial frequency: low or high).

Using a LOOCV procedure, we obtained unbiased single-trial responses of the classifiers. We then grouped these single-trial estimates into five bins relative to occipital theta, alpha and beta power, and contrasted the decoding accuracy between bins of the lowest and highest pre-cue power. In accordance with our alpha predictions, we found significant differences contrasting decoding accuracy between bins of low and high pre-stimulus occipital alpha power for pre-cue instruction (Figure 8A; p = 0.024; t = 0.2 to 0.21 and 0.47 to 0.49 s). In other words, decoding accuracy was higher for cues that were preceded by low alpha power. No significant effects were found for the decodability of retro-cue instruction by alpha power (p = 0.37).

**Figure 8.**
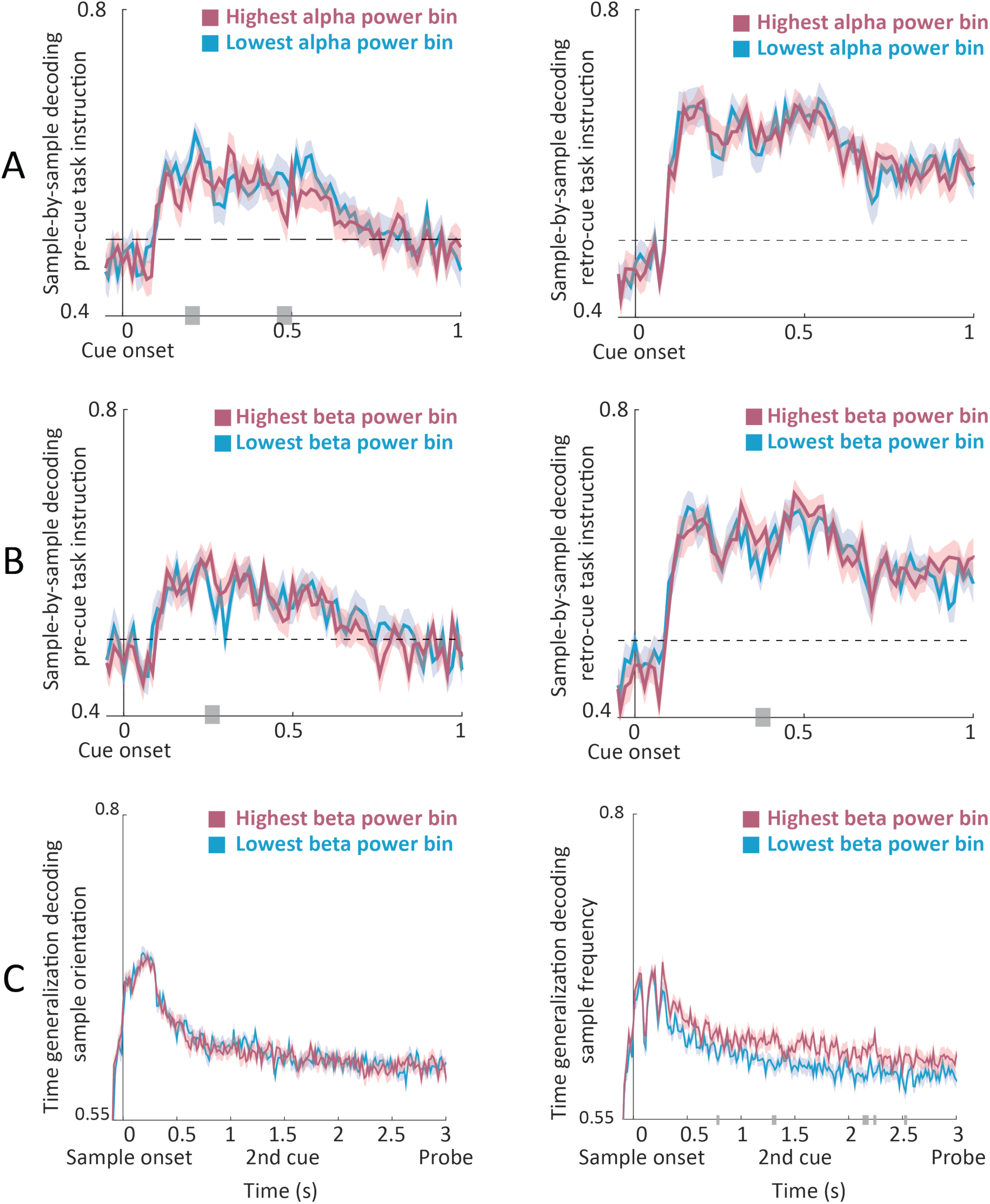
Effects of pre-stimulus oscillatory power on decodability. [A] Temporal sample-by-sample decoding of pre-cue (left panel) and retro-cue (right panel) instruction for the bins of lowest and highest occipital alpha power preceding cue onset. [B] Same as A for the beta band. [C] Averaged time-generalization matrices (over the first 0.3 s of training time) of decoding the sample’s orientation (left panel) and frequency (right panel) for the bins of lowest and highest beta power preceding sample onset. Gray bars indicate significant differences between low and high power bins.

In accordance with our beta predictions, we found significant differences contrasting decoding accuracy between bins of low and high beta power for pre-cue instruction (Figure 8B; p = 0.048; t = 0.26 to 0.27 s). In addition, we found a trend contrasting decoding accuracy between bins of low and high beta power for retro-cue instruction (p = 0.057; t = 0.37 to 0.39 s). Thus, contrary to alpha power, decoding accuracy was higher for cues that were preceded by high beta power. No significant modulations were found for the decodability of pre- or retro-cue instruction by theta power (p > 0.5 and p = 0.21, respectively), suggesting that this modulation of neural stimulus information was specific to alpha and beta oscillations.

In order to further test our predictions regarding beta power and (reactivation of) information content, we focused on decoding the spatial frequency and orientation of the sample stimulus. To this end, we applied a time-generalization approach in which the classifier was trained at each time point and tested on all other time points, thus allowing us to test the temporal stability of the information content. This approach yields a two-dimensional matrix across training and testing times. In order to reduce dimensionality, we averaged decoding accuracy across the early training time points (0 to 0.3 s post sample onset). We then contrasted the averaged decoding accuracy between the bins of lowest and highest pre-stimulus power.

In line with our predictions, we found significant differences between the averaged time-generalization matrices for decoding spatial frequency of the sample stimulus between bins of low and high occipital beta power (Figure 8C; p = 0.022). In other words, in trials where the sample was preceded by high beta power, classifiers trained on early time points more accurately decoded the sample’s spatial frequency when tested on later time points (0.8 to 2.5 s) in comparison to trials preceded by low beta power. No significant effects were found for the decodability of the sample’s orientation (p = 0.47) based on beta binning, though note that overall decoding performance was much lower for orientation than frequency (compare Figure 3 panels C and D for overall decoding performance along the diagonal).

No significant effects were found for the decodability of the sample’s spatial frequency or orientation when binning was performed according to theta (p = 0.11 and p = 0.2, respectively) or alpha power (p = 0.38, p = 0.46), suggesting that the maintenance of neural stimulus information was specifically supported by beta oscillations.

In summary, alpha and beta oscillations differentially modulated the information content of neural activity. We found a negative relationship between alpha power and cue representations, as predicted by the inhibitory gating account, while we found a positive relationship between beta power and cue and sample representations, as predicted by the circuit recruitment account.

## 4 Discussion

In this study, we used a delayed match-to-sample working-memory task with pre- and retro-cueing in order to study a comprehensive framework that integrates the functional roles of specific neural oscillations, and, critically, the coordination between them. We directly tested our proposed framework in which slow rhythms sample task-relevant input, while alpha/beta gates the information flow by suppressing task-irrelevant processing, and beta further serves to recruit task-relevant circuits. Crucially, via phase-amplitude coupling, the timing of faster oscillations is controlled by slower rhythms. Indeed, in line with their role as a pace keeper of input sampling, we found that successful matching of probe and sample stimuli was preceded by an increased phase alignment that was confined to slow rhythms. As for the gate keeping role of alpha oscillations, we demonstrated a pattern of alpha decrease that temporally tracked the presentation of the informative cue, correlated with faster behavioral performance and increased neural information content (indexed by decoding performance). Importantly, we shed new light on the dual role of beta oscillations: on the one hand, an alpha-like inhibitory role, gating the processing of relevant input, and on the other hand, a circuit-activation role, relevant for (reactivation of) information maintenance.

### 4.1 Slow oscillations: the pace keeper

The first tenet of our framework is slow oscillations (1–6 Hz): we hypothesized that their phase regulates the sampling of task-relevant input and controls the timing of faster oscillations via phase-amplitude coupling. Indeed, we demonstrate that inter-trial coherence (i.e., phase alignment) of slow oscillations was more prominent after presentation of the informative cues, and was followed by faster responses. Interestingly, this effect occurred before probe onset and was maximal in the pre- and post-central gyri and lateral prefrontal cortex. We also found that on fast trials, the phase of slow oscillations was more strongly coupled to the amplitude of faster oscillations (in the high beta range) in parietal areas.

We posit that this behavioral facilitation relies on more optimal temporal coordination reflected in the phase adjustment of slow oscillations and their coupling to faster oscillations, which in turn reflects better temporal prediction of probe onset. This is in line with previous reports that temporal predictions are associated with improved behavioral performance (Coull & Nobre, 1998; Miniussi et al., 1999; Nobre et al., 2007) and increased phase alignment of slow oscillations (Breska & Deouell, 2017), as well as recent evidence that the phase of slow oscillations predicts behavioral performance (Herbst & Obleser, 2019). This finding further corroborates previous reports of cross-frequency interactions during working memory (Bahramisharif et al., 2018; Bastos et al., 2018; Berger et al., 2019) and adds to a growing number of studies demonstrating the behavioral relevance of such coupling using intracranial (Axmacher et al., 2010) and scalp (Puszta et al., 2020) EEG. Thus, temporal coordination by slow oscillations governs the sampling of incoming stimuli (Busch et al., 2009; Fiebelkorn et al., 2013; Landau & Fries, 2012; VanRullen et al., 2011) and controls the timing of faster oscillations.

Finally, in addition to these delta-to-theta dynamics, we found that faster behavioral performance (and presentation of informative cues) cooccurred with stronger ERFs preceding and following probe onset (i.e., contingent negative variation or CNV, and N1/P2 and P3, respectively). This ERF enhancement may reflect better temporal prediction of probe onset in fast trials, resulting in a more prominent anticipatory ramping up of neural activity (Bidet-Caulet et al., 2015; Breska & Deouell, 2017; Brunia & van Boxtel, 2001). One potential concern is that our spectral analysis picked up on this evoked activity (rather than genuine oscillatory dynamics). However, given that our phase results were not substantially altered by removing evoked activity and that delta-to-theta and ERF dynamics were not identically localized, we posit that these changes in evoked activity reflect an additional aspect of anticipatory mechanisms that are boosted by slow rhythm phase modulations, i.e., enhanced phase alignment leads to a stronger evoked response.

In summary, we outline how slow oscillations not only coordinate input sampling but also faster oscillatory dynamics for optimal information processing and subsequent performance. Future research should investigate how this endogenous rhythmic sampling mechanism is deployed in contexts of fixed versus varying temporal structure.

### 4.2 Alpha oscillations: the gate keeper of information processing

We found that decreases in alpha power actively track the presentation of informative cues; i.e., regardless of its position in time, the informative cue was followed by a more prominent decrease in occipitoparietal alpha power. Our results are in line with the proposed gating role for alpha oscillations: once an input has been sampled through slow oscillations, its subsequent processing depends on its task relevance, i.e., alpha oscillations up- and down-regulate cortical excitability in task-relevant and irrelevant networks, respectively (Jensen & Mazaheri, 2010; Klimesch et al., 2007). This regulation of cortical excitability has been shown to occur spontaneously (van Dijk et al., 2008; Iemi et al., 2021; Linkenkaer-Hansen et al., 2004; Samaha et al., 2017) or to track relevant input in space (ElShafei et al., 2018; Haegens et al., 2012; Mazaheri et al., 2014; Thut et al., 2006) or, as in the case of the current study, in time (Hanslmayr et al., 2011; Rohenkohl & Nobre, 2011; van Diepen et al., 2015; van Ede et al., 2017).

In addition, we found that faster responses were preceded by more prominent occipitoparietal alpha power decrease. This is in line with previous studies demonstrating that behavioral performance correlates negatively with alpha activity in task-relevant (Jiang et al., 2015; van Ede et al., 2017) and positively in task-irrelevant regions (Haegens et al., 2010; Yu et al., 2017). Both lines of results reflect the same low-level gating mechanism deployed to allocate more resources to task-relevant networks, either by directly boosting the processing of task-relevant information or by suppressing that of task-irrelevant information (i.e., distracting or competing stimuli).

Importantly, we demonstrate that pre-stimulus alpha power decrease enhances neural decodability (i.e., sensory representations) of subsequent task-relevant items. These results are in line with previous work reporting a negative relationship between decoding performance and pre-stimulus alpha power (Barne et al., 2020; van Ede et al., 2018), and shows how alpha through regulating cortical excitability might influence sensory representations.

### 4.3 Beta oscillations: a dual role in information gating and circuit formation

We hypothesized that multiple beta mechanisms exist, with one beta mechanism playing a local functionally inhibitory role (“inhibitory beta”), similar to alpha, and another beta mechanism allowing top-down flexible activation of task-relevant circuits that maintain the information required to successfully perform the task at hand (“circuit beta”). These two mechanisms translate to opposite power modulations.

In support for its inhibitory role, we show that similarly to alpha, decreases in prefrontal and occipitoparietal beta power are contingent upon the occurrence of the informative cue, and associated with faster responses. This is in line with previous reports on the alpha-like gating role played by beta oscillations (Bastos et al., 2018; van Ede et al., 2011; Miller et al., 2018).

Interestingly, while there is overlap in functionality and temporal patterns between alpha and beta, the localization of the inhibitory beta effects extends more towards frontoparietal regions, relative to that of the alpha effects.

In support for its circuit role, we demonstrate that increases (rather than decreases) in beta power index a stronger decoding and maintenance of task-relevant features, i.e., the decodability of stimuli preceded by high beta power was boosted and generalized for longer durations. This is in line with previous studies demonstrating that beta power correlates with stimulus information maintained in working memory as well as subsequent decision outcomes (Haegens et al., 2017; Proskovec et al., 2019; von Lautz et al., 2017). Here we go one step further, by directly linking the power of beta oscillations to the quality of information content in neural activity patterns. We posit that stronger beta synchronization indexes a stronger (longer lasting) setting up of task-relevant circuits (i.e., working-memory nodes). In turn, this enhanced circuit recruitment leads to more efficient maintenance of stimulus features and thus allows them to be decoded for longer durations.

Taken together, our results corroborate the proposed dual roles for beta rhythms in our framework (Miller et al., 2018; Spitzer & Haegens, 2017). Future invasive studies are required to dissociate the underlying generators (e.g., on the cell circuit/laminar level) of these proposed dual betas. In addition, the temporal and spatial evolution of alpha and beta rhythms should be thoroughly compared, in order to highlight potential differences and similarities in their roles in active inhibition.

### 4.4 Conclusion

In summary, we demonstrate how rhythms from the delta to beta range might subserve higher-level cognitive functions by providing low-level mechanistic operations. Combined, these oscillatory building blocks allow for selective sampling of input, disengaging and engaging task-irrelevant and relevant networks, and temporal organization of these respective dynamics. Critically, we find that these oscillatory dynamics correlate both with the behavioral performance of the participant, and with the information content in the recorded brain signal.

## Author Contributions

HE, YZ and SH Designed Research; HE and YZ Performed Research; HE and YZ Analyzed data; HE, YZ and SH Wrote the paper

## Acknowledgements

We would like to thank Kristina Baumgart for her assistance with data acquisition, and Luca Iemi for thoughtful comments on an earlier version of the manuscript.

